# Characteristics and Evolutionary analysis of Photosynthetic Gene Clusters on Extrachromosomal Replicons: from streamlined plasmids to chromids

**DOI:** 10.1101/663864

**Authors:** Yanting Liu, Qiang Zheng, Wenxin Lin, Nianzhi Jiao

## Abstract

Aerobic anoxygenic photoheterotrophic bacteria (AAPB) represent intermediates in the evolution from photoautotrophic to heterotrophic metabolisms. Substantial evidence indicates that highly conserved photosynthetic gene clusters (PGCs) of AAPB can be transferred between species, genera, and even phyla. Furthermore, analysis of recently discovered PGCs carried by extrachromosomal replicons (exPGCs) suggests that extrachromosomal replicons (ECRs) play an important role in the transfer of PGCs. In the present study, thirteen *Roseobacter* clade genomes from seven genera that harbored exPGCs were used to analyze characteristics and evolution of PGCs. The identification of plasmid-like and chromid-like ECRs from PGC-containing ECRs revealed two different functions: the spread of PGCs among strains and the maintenance of PGCs within genomes. Phylogenetic analyses indicated two independent origins of exPGCs, corresponding to PufC-containing and PufX-containing photosynthetic reaction complexes. Furthermore, the two different types of complexes were observed within different strains of the same *Tateyamaria* and *Jannaschia* genera. The two different complexes were also differentially carried by chromosomes and ECRs in the strains, respectively, which provided clear evidence for ECR-mediated PGC transfer. Multiple recombination events of exPGCs were also observed, wherein the majority of exPGCs were inserted by replication modules at the same genomic positions. However, the exPGCs of the *Jannaschia* strains comprised superoperons without evidence of insertion, and therefore likely represent an initial evolutionary stage where the PGC was translocated from chromosomes to ECRs without further combinations. Lastly, a scenario of PGC gain and loss is proposed that specifically focuses on ECR-mediated exPGC transfer to explain the evolution and patchy distribution of AAPB within the *Roseobacter* clade.

**Importance:** The evolution of photosynthesis was a significant event during the diversification of biological life. Aerobic anoxygenic heterotrophic bacteria (AAPB) share physiological characteristics with both photoautotrophs and heterotrophs and are therefore suggested to be evolutionary intermediates between the two lifestyles. Here, characterization and evolutionary analyses were conducted for thirteen bacterial strains that contained photosynthetic gene clusters (PGCs) carried by extrachromosomal replicons (ECRs) to shed light on the evolution of photosynthesis in bacteria. Specifically, these analyses improved the “Think Pink” scenario of PGC transfer that is mediated by ECRs in *Roseobacter* clade strains. This study advances our understanding of the importance of ECRs in the transfer of PGCs within marine photoheterotrophic bacteria.

## Introduction

Aerobic anoxygenic photoheterotrophic bacteria (AAPB) are photosynthetic members of the Proteobacteria phylum that contain bacteriochlorophyll *a* and are widely distributed in euphotic ocean environments (Kolber *et al.*, 2001; Jiao *et al.*, 2007; Yutin *et al.*, 2007; Koblížek *et al.*, 2007; Jiao *et al.*, 2009; Ritchie and Johnson, 2012; Koblížek, 2015). AAPB account for up to 10% of the total bacterial communities in the upper ocean layers and play crucial roles in carbon and energy cycling (Cottrell *et al.*, 2006; Lami *et al.*, 2007; Boeuf *et al.*, 2013). AAPB are facultatively phototrophic, using light as an alternative energy source, which thereby reduces the respiration of organic carbon from marine primary production by ~2.4–5.4% (Jiao *et al.*, 2009). In addition to their ecological significance, physiological and genomic analysis of AAPB have indicated their important potential to inform on the evolution of photosynthesis (Yurkov and Csotonyi, 2009a; Yurkov and Hughes, 2017; Beatty, 2002). AAPB are hypothesized to have evolved from anaerobic phototrophs after widespread oxygenation of Earth’s atmosphere ~2.4-2.3Ga ago (Bekker *et al.*, 2004; Holland, 2006). Purple photosynthetic bacteria are one such group of anaerobic phototrophs that are thought to represent the ancestral lineage to AAPB due to the high similarity of photosynthetic apparatuses between them (Yurkov and Beatty, 1998; Shimada, 2004; Yurkov and Csotonyi, 2009). However, unlike purple photosynthetic bacteria that grow in the absence of oxygen and are mainly autotrophic, AAPB are aerobic heterotrophs that perform photosynthesis as an auxiliary energy conservation strategy, which can contribute up to 20% of their total cellular metabolic energy demands (Yurkov and Beatty, 1998; Kolber *et al.*, 2001). Phylogenetic analysis of 16S rRNA gene indicates that AAPB are closely related to non-phototrophic bacteria and purple non-sulfur bacteria within the Proteobacteria (Nishimura *et al.*, 1996; Yurkov and Beatty, 1998; Beatty, 2002). Consequently, it has been proposed that AAPB could represent intermediates in the evolution of heterotrophs from photoautotrophs (Nishimura et al., 1996; Yurkov and Beatty, 1998; Keppen *et al.*, 2013; Thiel *et al.*, 2018).

Similar to purple photosynthetic bacteria, photosynthetic gene clusters (PGCs) are found in AAPB. The large PGC superoperon is approximately 35–50 kb in length and contains approximately 40 genes required for the biosynthesis of bacteriochlorophyll, carotenoids, photosynthetic reaction complexes, and light harvesting complexes, in addition to other regulatory functions (Beatty, 1995; Liotenberg *et al.*, 2008; Swingley *et al.*, 2009; Zheng *et al.*, 2011). PGC gene sequences are highly conserved and some essential genes within PGCs (e.g., *pufL, pufM,* and *bchY)* in whole PGCs are typically used as molecular markers in phylogenetic analyses to classify phototrophic Proteobacteria and study their ecological diversity and evolutionary relationships (Beja *et al.*, 1992; Yutin *et al.*, 2007; Zeng *et al.*, 2014; Zheng *et al.*, 2016).

Previous comparisons of closely related strains have demonstrated that PGCs can be lost from bacterial genomes. For example, *Citromicrobium* sp. JL1363 lost PGCs from its genome during its evolutionary history and is now reliant solely on heterotrophy (Zheng *et al.*, 2012). Furthermore, PGCs can be transferred between proteobacterial strains and among distantly related phyla. For instance, although *Erythrobacter* sp. AP23 shares 99.5% 16S rRNA gene sequence identity with *Erythrobacter* sp. LAMA915, only the former strain contains a PGC within its genome (Zheng *et al.*, 2016). Intriguingly, the PGC of *Erythrobacter* sp. AP23 is closely related to that of *Citromicrobium* (Zheng *et al.*, 2016). Moreover, the Gemmatimonadetes strain AP64 is a novel organism that contains bacteriochlorophyll *a* (BChl a), and has been suggested to have acquired its PGC from photoheterotrophic bacteria of the Proteobacteria via an ancient horizontal gene transfer (HGT) event (Zeng *et al.*, 2014).

Members of the *Roseobacter* clade are important ecological generalists within marine ecosystems and many are AAPB (Brinkhoff *et al.*, 2008; Newton *et al.*, 2010; Luo and Moran, 2014). Inconsistencies within phylogenetic topologies between the 16S rRNA gene and PGC indicate that HGT events of PGCs have occurred among members of this clade (Nagashima *et al.*, 1997; Raymond, 2002; Petersen *et al.*, 2012; Brinkmann *et al.*, 2018). Of note, PGC-containing extrachromosomal replicons (ECRs) have been identified within six strains of the *Roseobacter* clade, further implicating the possibility of HGT of PGCs among these species (Brinkmann *et al.*, 2018). ECRs are mobile genetic elements that carry many essential genes involved in metabolism that are essential for rapid adaptation to changing environments (Sørensen *et al.*, 2005; Thomas and Nielsen, 2005; Petersen *et al.*, 2013). Therefore, it is likely that PGCs can also be transferred via ECRs. However, limited study have been focused on the role of ECRs in the HGT of PGCs (Petersen *et al.*, 2012; Brinkmann *et al.*, 2018).

In this study, we analyzed thirteen *Roseobacter* clade genomes that contain PGCs carried by extrachromosomal replicons (exPGCs). Phylogenetic and structural analysis of the exPGCs, in addition to comparisions against chromosomal PGCs (cPGC), were used to elucidate the potential role of ECRs in the transfer of PGCs among *Roseobacter* clade species.

## Results and Discussion

### General features of the thirteen *Roseobacter* clade strains

Thirteen *Roseobacter* clade strains that carried PGC-containing ECRs and were affiliated with seven different genera: *Tateyamaria, Jannaschia, Sulfitobacter, Roseobacter, Oceanicola, Shimia,* and *Nereida* (Table 1). Genome sizes of the thirteen strains ranged from 2.89 to 4.75 Mb, while the average genomic GC content ranged from 54.0 to 65.5%. Most strains contained more than two ECRs and the largest ECR among the thirteen strains was a 185 kb PGC-containing ECR in *Sulfitobacter* sp. AM1-D1. The thirteen identified exPGCs ranged in size from 41.3 to 54.0 kb and their average GC contents were consistent with their respective chromosomes. All PGC-containing ECRs harbored DnaA-like I replication systems except for that of *S. guttiformis,* which exhibited a RepB-III type system.

**Table 1.**
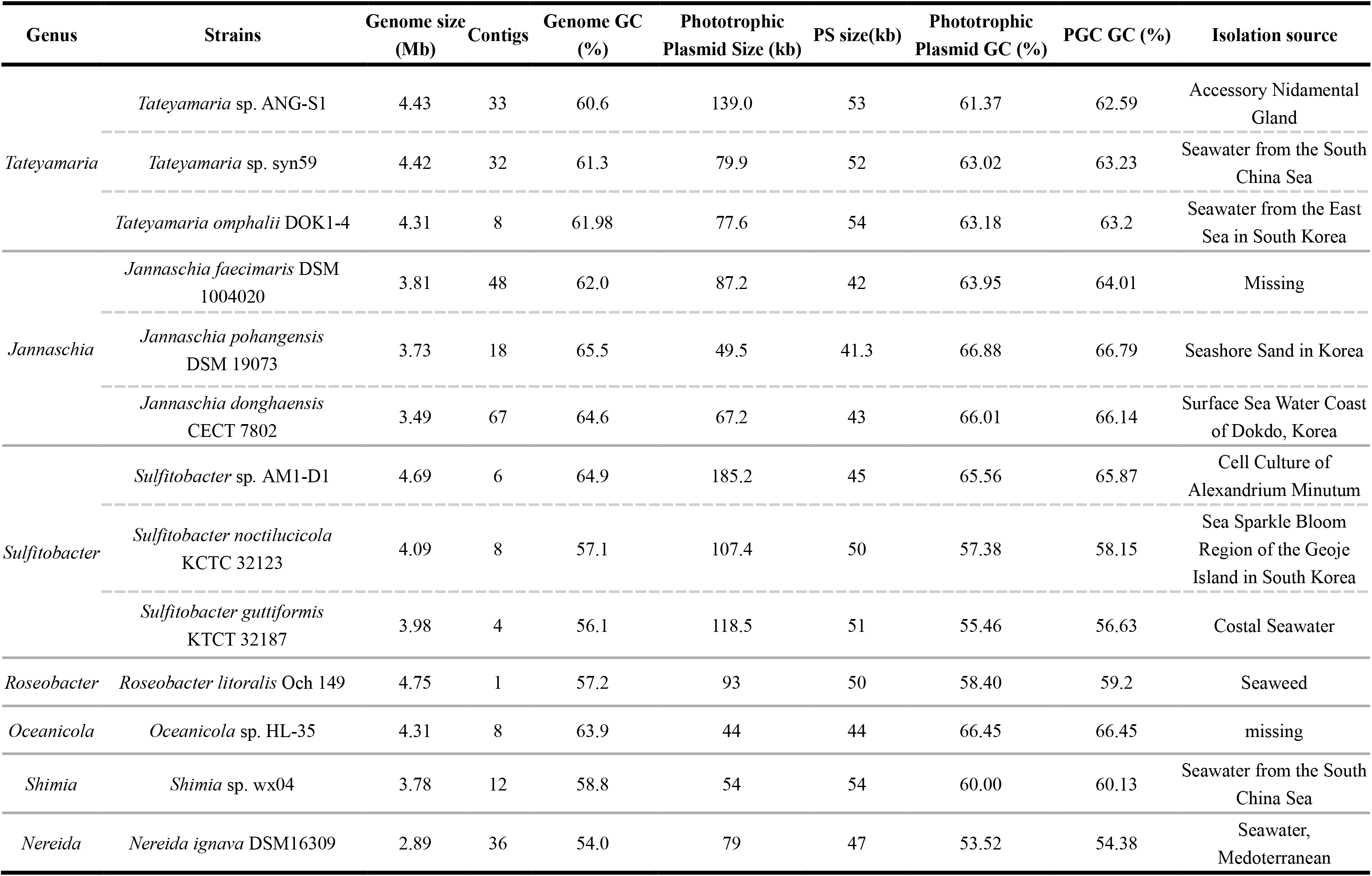
Genomic information for thirteen *Roseobacter* strains carrying exPGCs.

*Tateyamaria* have been isolated from different environments including seawater, tidal-flat sediments, and marine animals (Kurahashi and Yokota, 2007; Sass *et al.*, 2010; Jeanthon *et al.*, 2011; Collins *et al.*, 2015; Reis and Costa, 2018). *Tateyamaria* have also been frequently detected in algal culture bacterial communities and can occasionally dominate such communities (Bengtsson *et al.*, 2012; Kalitnik *et al.*, 2017; Huggett *et al.*, 2018). The *pufM* gene is present in all genomes reported for this genus (as of 2/28/2019). Three strains carrying exPGCs with similar genome sizes (avg. 4.4 ± 0.8 Mb) and GC contents (avg. 61.30 ± 0.70%) were chosen for analysis in this study. The size of the ECRs carrying exPGCs ranged from 77.6 to 139.8 kb, and the exPGCs had similar sizes (avg. 53.0 ± 1.0 kb). A considerable number of highly homologous genes were present in the three PGC-containing ECRs of this genus (Fig. 1). In addition, genes present on smaller PGC-containing ECRs were mostly present on larger PGC-containing ECRs. Phylogenetic analysis of the replication partitioning gene (*parA*) indicated that the replication modules of the three PGC-containing ECRs were highly conserved, which was supported by high bootstrap values among homologs (Fig. S1).

**Fig. 1.**
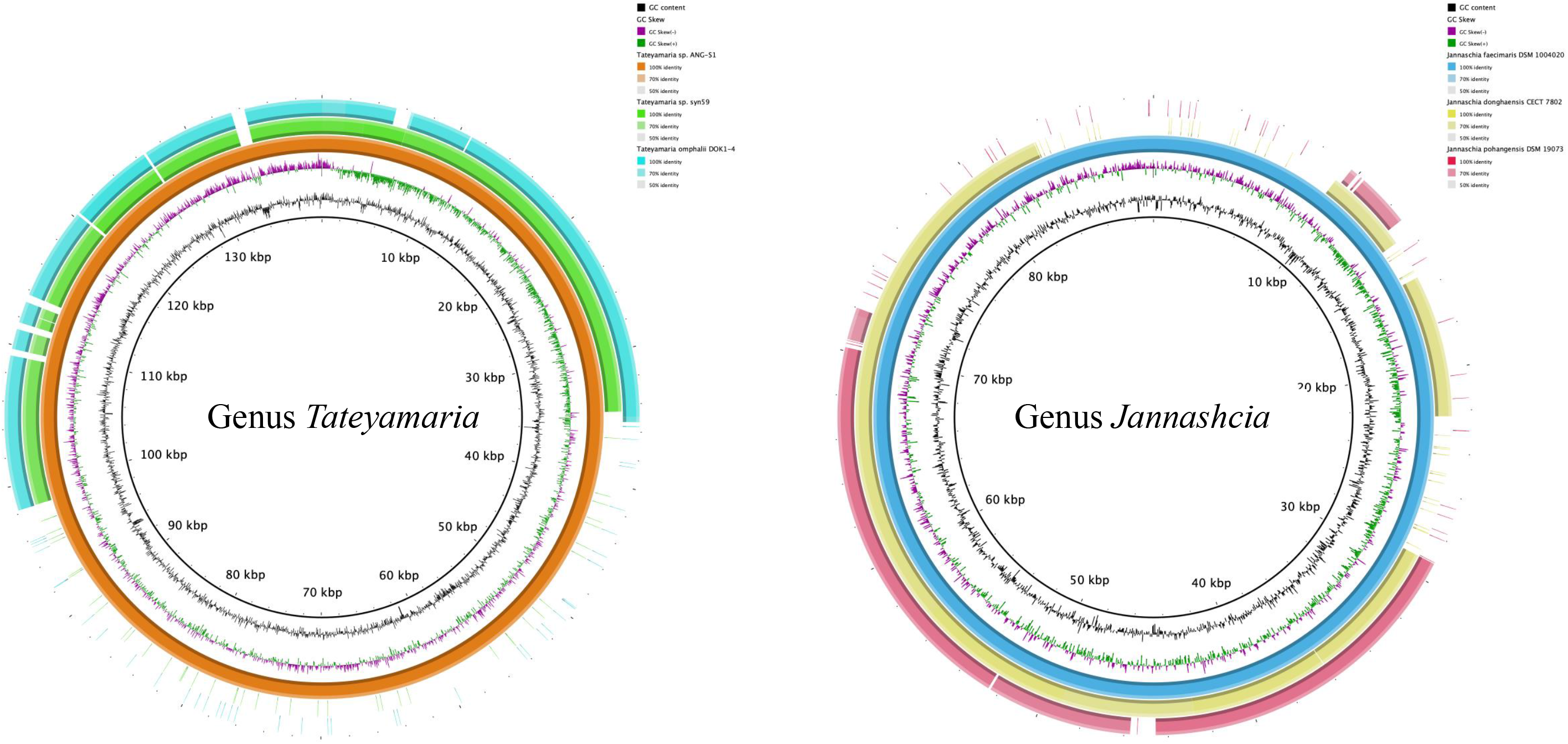
Circular maps of the PGC-containing extrachromosomal replicons for *Tateyamaria* (a) and *Jannaschia* (b) strains. (a) *Tateyamaria* sp. ANG-S1 is used as the reference. Tracks (moving outward) show G+C content, GC skew (G-C/G+C), *Tateyamaria* sp. ANG-S1 synteny, *Tateyamaria* sp. syn59 synteny, and *Tateyamaria* omphalii DOK1-4 synteny. (b) *Jannaschia faecimaris* DSM 1004020 is used as the reference. Tracks (moving outwards) show: G+C content, GC skew (G-C/G+C), *Jannaschia faecimaris* DSM 1004020 synteny, *Jannaschia donghaensis* CECT 7802 synteny, and *Jannaschia pohangensis* DSM19073 synteny.

*Jannaschia* is an ecologically important genus of AAPB (Zeng *et al.*, 2007). Indeed, 25-30% of all AAPB 16S rRNA gene clone sequences in Central Baltic Sea belonged to *Jannaschia-related* bacteria (Salka *et al.*, 2008). Among known *Jannaschia* isolates, CCS1 is the only strain observed to conduct photoheterotrophy (Moran *et al.*, 2007; Pujalte *et al.*, 2014). Twelve *Jannaschia* strains were available for analysis with whole genome sequences, with six containing PGCs, wherein three were exPGC types and the others were cPGC types. The PGC-containing ECRs ranged in size from 49.5 to 87.2 kb, while their exPGCs were ~45 kb in length. The complete genomes of the three *Jannaschia* strains carrying exPGCs ranged from 3.49 to 3.81 Mb, with GC contents ranging from 62.0 to 65.5%. The three PGC-containing ECRs of *Jannaschia* shared a large syntenic region comprising ~50 kb and the larger PGC-containing ECRs contained genes that were carried by the PGC-containing smaller ECRs, as observed for the ECRs in the *Tateyarima* genomes (Fig. 1). In addition, their replicon replication modules were closely related based on a phylogenetic analysis of *parA* genes (Fig. S1).

*Sulfitobacter* are widely distributed in different marine environments and may play important roles in organic sulfur cycling (Pukall *et al.*, 1999; Labrenz *et al.*, 2000; Rooney-Varga *et al.*, 2005; Prabagaran *et al.*, 2007; Moran *et al.*, 2010; Zachariah *et al.*, 2017). Culture-independent surveys of AAPBs have indicated that *Sulfitobacter* account for a significant fraction of AAPB communities in natural environments (Buchan *et al.*, 2005; Boeuf *et al.*, 2013; Yinxin Zeng *et al.*, 2016). However, only a few *Sulfitobacter* AAPB strains have been isolated and described (Labrenz *et al.*, 2000; Boeuf *et al.*, 2013; Kwak *et al.*, 2014). Genomes from only three *Sulfitobacter* AAPB strains have been sequenced, which has revealed that their PGCs were located on ECRs. The three genomes ranged in size from 3.98 to 4.69 Mb, with GC contents varying from 56.1 to 64.9%. The three PGC-containing ECRs exhibited sizes greater than 100 kb, while the three exPGCs were 45, 50, and 51 kb. Other than genes involved in photosynthesis, only a small number of genes were shared by PGC-containing ECRs in *Sulfitobacter,* which contrasted with the high degree of conservation observed for ECRs of *Tateyamaria* and *Jannaschia.* Furthermore, the *parA* genes within the three ECRs of *Sulfitobacter* were phylogenetically very distinct (Fig. S1).

*Roseobacter litoralis* Och 149 was the first AAPB bacterium described in 1991, and contained a linear ECR harboring a PGC (Shiba, 1991; Kalhoefer *et al.*, 2011). The genus only comprises two species, *R. litoralis* Och 149 and *R. denitrificans* OCh114, with the latter containing a cPGC (Shiba, 1991). In contrast, little is known about phototrophic species within *Oceanicola, Shimia,* and *Nereida. Oceanicola* sp. HL-35 exhibits 99% 16S rRNA gene sequence identity with *Lacimonas salitoletrans* TS-T30, which lacks a PGC (Zhong *et al.*, 2015). Prior to the isolation of *Shimia* sp. wx04, *Shimia* were thought to be strict chemoorganotrophs based upon culture-dependent investigations (Choi and Cho, 2006; Chen *et al.*, 2011; Hameed *et al.*, 2013; Hyun *et al.*, 2013; Nogi *et al.*, 2015). Lastly, *N. ignava* DSM 16093 was isolated from waters of the Mediterranean and is the sole species currently described for the *Nereida* (Pujalte *et al.*, 2005; Arahal *et al.*, 2016).

### Identification of PGC-containing chromids and PGC-containing plasmids

Chromids are a novel type of ECR that were recently described as distinct genetic elements from both chromosomes and plasmids (Harrison *et al.*, 2010). Since chromids typically carry essential genes, they more stably remain than plasmids in bacteria and are considered indispensable for bacterial hosts (Harrison *et al.*, 2010). The comparison of relative synonymous codon usage (RSCU) of bacterial chromosomes and ECRs can help identify if ECRs are chromids or otherwise, as RSCU of chromid are similar to those of corresponding chromosomes (Harrison *et al.*, 2010; Petersen *et al.*, 2013; Petersen and Wagner-Döbler, 2017). Principal component analysis (PCoA) of the RSCU of all replicons from the thirteen strains were analyzed to be used to the classification of elements as chromids and plasmids. The analyses indicated that seven PGC-containing ECRs *(Tateyamaria* sp. ANG-S1, *Tateyamaria* sp. syn59, *Tateyamaria omphalii* DOK1-4, *Sulfitobacter* sp. AM1-D1, *Sulfitobacter noctilucicola* KCTC 32123, *Sulfitobacter guttiformis* KTCT 32187, and *Roseobacter litoralis* Och 149, *Nereida ignava* DSM16309) could be clearly assigned as chromid-like ECRs, while the other ECRs containing PGCs were provisionally classified as plasmid-like ECRs (Fig. 2).

**Fig. 2.**
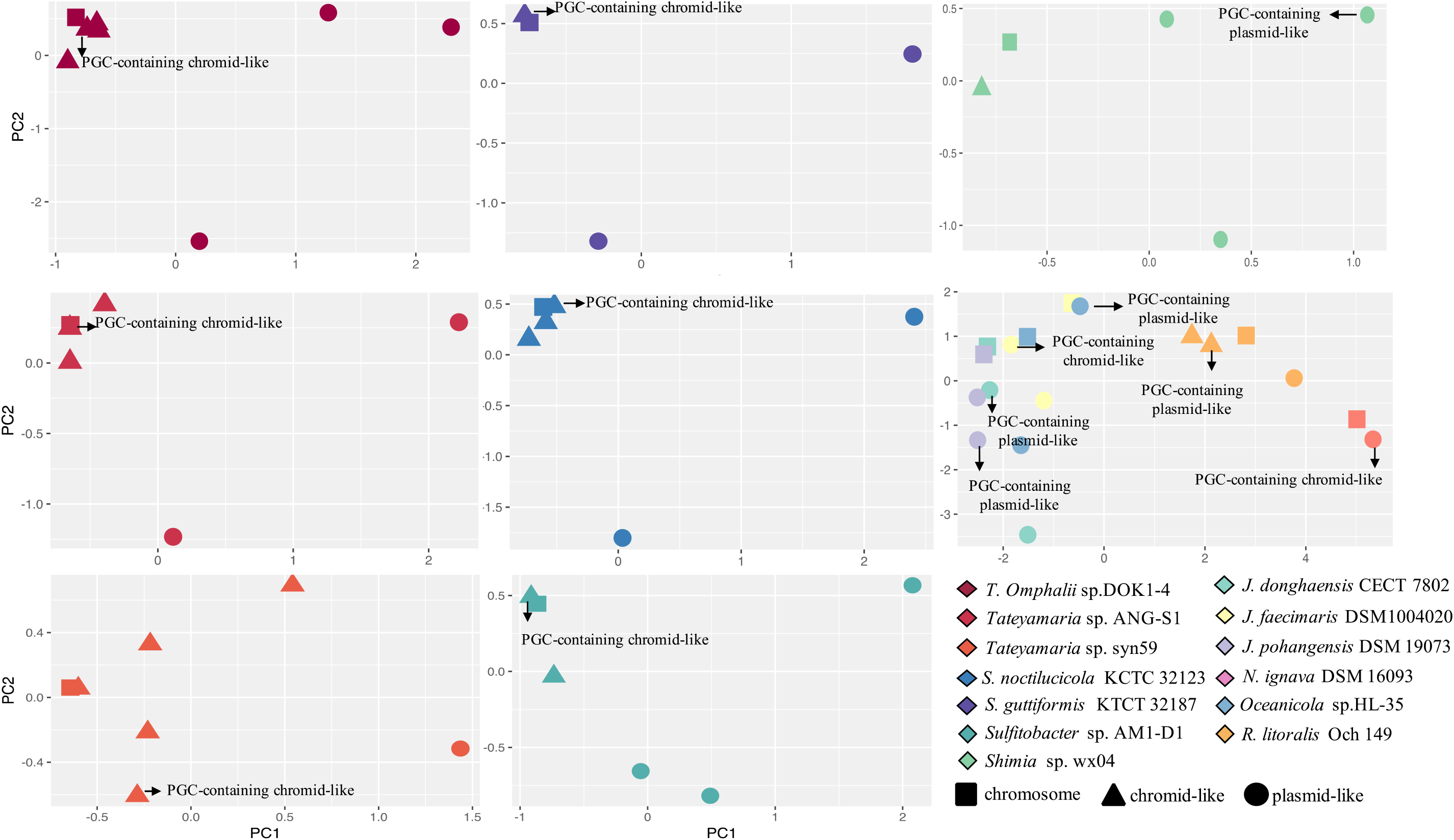
Principal component analysis of the relative synonymous codon usage (RSCU) of replicons from the thirteen strains carrying exPGCs. Chromsomes, chromids, and plasmids are indicated by squares, triangles, and circles, respectively. The strains containing greater than three replicons were analyzed by strain, while the rest were evaluated using a cross-strain analysis.

### Phylogenetic analysis

A phylogenetic analysis based on 16S rRNA gene nucleotide sequences from thirteen exPGC-containing bacteria and 43 reference strains was conducted with photoheterotrophs and heterotrophs to show the phylogenetic distribution of the thirteen strains carrying exPGCs (Fig. S2). As observed for the cPGC-containing AAPB, the thirteen exPGC-containing bacterial strains did not comprise a monophyletic phylogenetic group, but were instead distributed throughout the 16S rRNA phylogenetic tree.

To further investigate the phylogenetic relationships of the *Roseobacter* clade strains, 38 photoheterotrophic bacterial genomes were subjected to phylogenetic analysis using 29 conserved PGC genes. Comparison of phylogenies based on 16S rRNA nucleotide sequences and amino acid sequences of 29 conserved genes of the PGC revealed considerable topological differences (Fig. 3). For example, the four *Tateyamaria* strains clustered and closely related to the *N. ignava* DSM16309 strain in the 16S rRNA gene phylogenetic analysis. However, the *Tateyamaria* strains were using the highly conserved 29-gene of the PGC, with *Tateyamaria.* sp. Alg231-49 closely related to *Thalassonium* sp. R2A62 and the other three strains associated with the *Roseobacter* species. Similarly, the five *Jannaschia* strains formed a group with bootstrap support (86%) based on the 16S rRNA phylogenetic analysis; however, they were clearly differentiated into two distant subtrees in the PGC-based phylogenetic analysis.

**Fig. 3.**
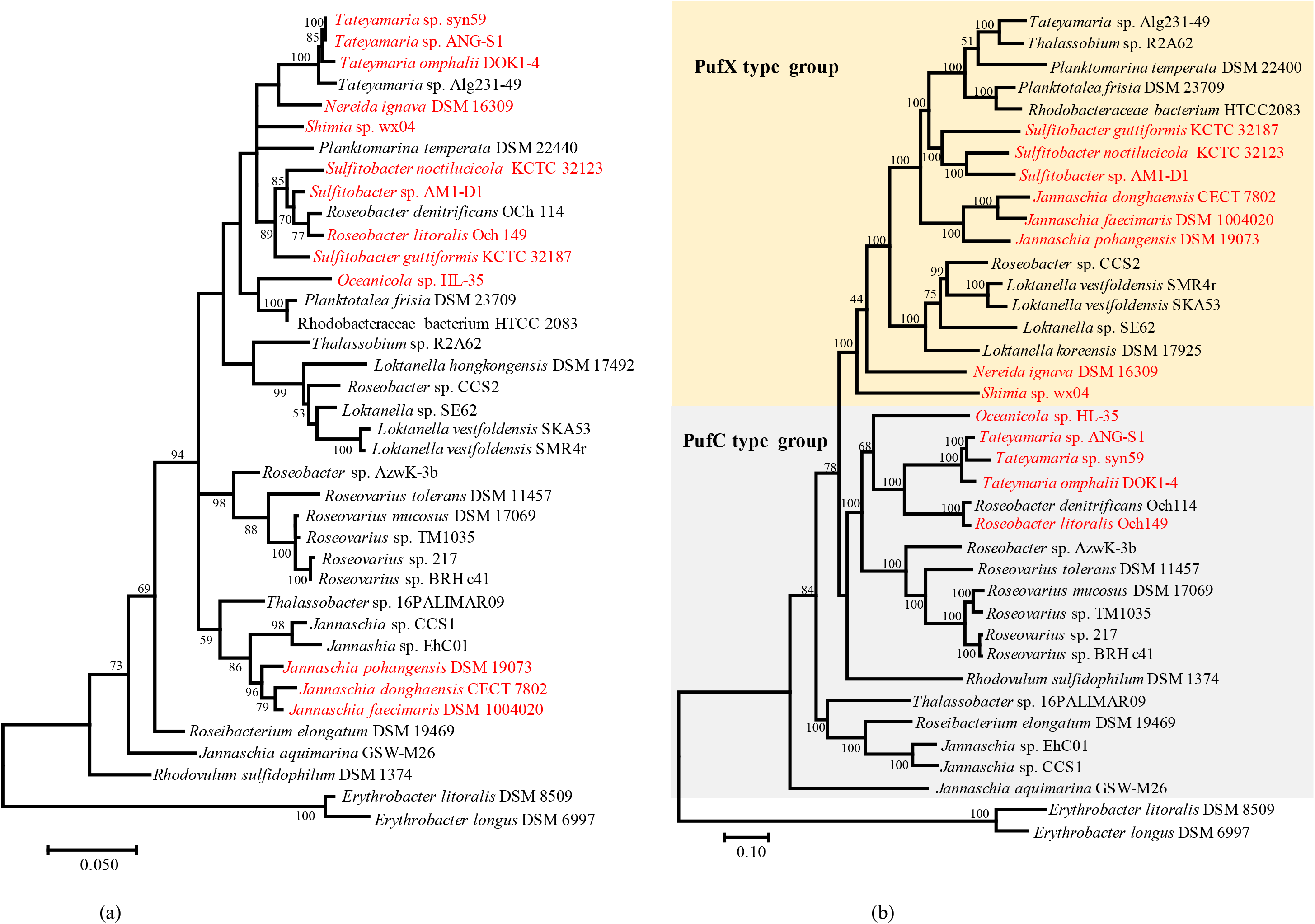
Phylogenetic trees of 16S rRNA genes (a) and 29 conserved PGC genes concatenation (b). Trees were constructed using maximum likelihood methods with 100 bootstrap replicates to evaluate node support. Only bootstrap values > 50% are shown. Phototrophic strains carrying exPGCs are indicated in red. Accession numbers of all sequences are summarized in Table S1.

In addition, the PGC-based phylogenetic analysis indicated the presence of two phylogenetic groups corresponding to differences in photosynthetic reaction complex types. Specifically, the groups corresponded to PufC-containing and PufX-containing groups (Fig. 3b). In particular, *Tateyamaria* sp. syn59, *T. omphalli* pDOK1-4, *Tateyamaria* sp. ANG-S1, *R. litoralis* Och149, and *Oceanicola* sp. HL-35 contained *pufC* genes, while the others contained *pufX* genes. Five of the exPGCs in the PufC group clustered together, suggesting common ancestry for these exPGCs. Moreover, the cPGC in *Roseobacter denitrificans* Och114 represented a basal clade to a subtree also comprising the five aforementioned exPGCs, thereby providing evidence for chromosomal reintegration from an ECR (Brinkmann *et al.*, 2018). Furthermore, the externally nested position of the exPGC from *Oceanicola* sp. HL-35 within this subtree indicated that the exPGC was possibly transferred from other AAPB strains and could be further transferred to other distant bacterial strains via an ECR. Nine of the thirteen exPGCs belonged to three genera among the seven that were analyzed. Among these, the exPGCs from strains of the same genus were closely related phylogenetically, suggesting that the transfer of exPGCs was more likely to occur among strains within the same genus. Two photosynthetic reaction complex types were observed within the genomes of different strains within the *Tateyamaria* and *Jannaschia* genera. Among the six PGC-containing *Jannaschia* strains, three exPGCs were PufX types while the other three were PufC types. Similarly, among the four phototrophic *Tateyamaria* strains analyzed, one cPGC was a PufX type, while the other three exPGCs were PufC types.

### exPGC structures and arrangements

Three different structures were observed among the thirteen exPGCs (Fig. 4). The 41 photosynthetic genes on the ECR of the three *Jannaschia* strains were organized into one superoperon with the same structure as that of the cPGC (Zheng *et al.*, 2011). In addition, the exPGC of *Sulfitobacter* sp. AM1-D1 was separated by more than 100 genes between *bchIDO* and *hemECA.* The other exPGCs all appeared to have been inserted by their replication modules, as previously identified (Petersen *et al.*, 2012; Brinkmann *et al.*, 2018).

**Fig. 4.**
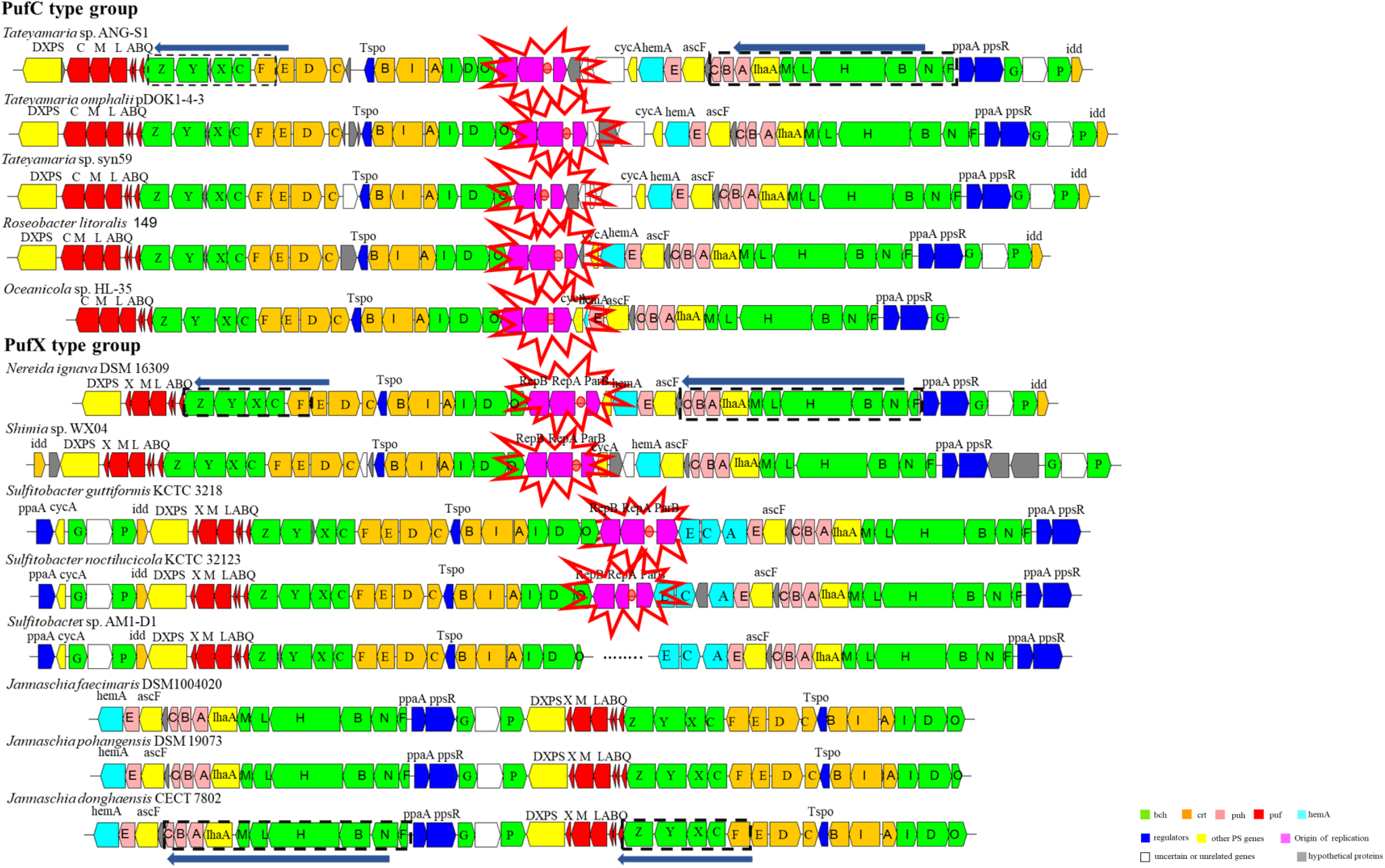
Photosynthetic gene cluster structures and arrangements for thirteen exPGCs. ECR module positions within the exPGC are shown in red and highlighted by stars.

The arrangement of photosynthetic genes is also a key characteristic of PGCs in AAPB. Three forms of PGC arrangement have been observed in *Roseobacter* clade organisms based on the combination of two conserved regions *(puh-LhaA-bchMLHNF* and *puf-bchZYXC-crtF)* (Zheng *et al.*, 2011). The thirteen exPGCs contained the same conserved gene order as cPGCs and comprised two different types. The type I arrangement was exhibited by exPGCs of the three *Jannaschia* strains *(J. pohangensis* DSM19073, *J. donghaensis* CECT7802, and *J. faecimaris* DSM10004020), wherein genes were arranged as forward *puh-LhaA-bchMLHNF* plus forward *puf-bchZYXC-crtF*. Type II arrangements were observed for the other 10 strains wherein arrangements followed the pattern of forward *puf-bchZYXC-crtF* plus forward *puh-LhaA-bchMLHNF* (Fig. 4).

The arrangement of exPGCs of the PufC-containing group were of type II, which exhibited the high conservation in the direction and order of all photosynthetic genes on the exPGCs. In contrast, the exPGCs in the PufX-containing group exhibited two different types of arrangements, with unique traits present in different genera. For example, *hemC* and *hemE* genes encoding tetrapyrrole biosynthesis proteins were only present in the exPGCs of *Sulfitobacter.* In addition, the genomic region ranging from *bchG* to *idd* was located upstream of the *puf* operon in *Sulfitobacter,* although it is typically downstream of the *ppaA* and *ppsR* regulator genes within PGCs (Zheng *et al.*, 2011). Furthermore, the three exPGCs of the *Jannaschia* strains lacked cytochrome c2 (cyc2) and diphosphate delta-isomerase (idi) genes. *cyc2* is involved in electron transfer while *idi* is involved in carotenoid biosynthesis. The loss of these genes is not lethal for phototrophic bacteria (Bonnett, 1995; Addlesee *et al.*, 2000), but the genes are nevertheless expected to be present in the PGCs of phototrophic bacteria within the *Roseobacter* clade (Jones *et al.*, 1990; Hahn *et al.*, 1996; Brinkmann *et al.*, 2018).

### Evidence of ECR-mediated PGC transfer within the *Roseobacter* clade

Recent studies have suggested that ECRs could be vehicles for HGT of PGCs, albeit with limited evidence (Petersen *et al.*, 2012; Brinkmann *et al.*, 2018). The transfer of PGCs by ECRs was supported in our analyses by the coexistence of two different types of photosynthetic reaction complexes (PufC and PufX types) in different strains of two genera, *Tateyamaria* and *Jannashcia*. In particular, these two types of photosynthetic reaction complexes were located on cPGCs and exPGCs, respectively. The Global Ocean Sampling expedition metageomes first revealed that *pufC* could be replaced by *pufX* in AAPB, and that *pufC* and *pufX* were present in different AAPB phylogroups (Yutin and Béjà, 2005; Yutin *et al.*, 2007). Thus, phylogenetic divergence of the two types of photosynthetic reaction complexes in strains from the same genus suggested that one or both of them were introduced by other phototrophic phylogroups. Moreover, phylogenetic congruence between whole PGCs and conserved photosynthetic operons within the PGC (i.e., *bchFNBHLM-IhaA-puhABC* and *pufMLABQ-bchZYXC-crtF)* indicate that PGCs act as entire functional units rather than being subject to partial transfer between strains (Fig. S3); this is consistent with a previous study (Brinkmann *et al.*, 2018). ECRs are mobile genetic elements and thus PGCs carried by ECRs are more likely to be horizontally transferred.

### The potential for transfer of PGC-containing plasmids

As described above, the thirteen PGC-containing ECRs were divided into two types based on their sizes and functions. Small PGC-containing ECRs within *Oceanicola* sp. HL-35, *Shimia* sp. wx04 and *J. pohangensis* DSM19073 carried more than 80% of the genes encoding for photosynthetic capacity. These ECRs are usually present as plasmids and are likely to play an important role in the transfer of photosynthetic capacity among species. This is especially probable because the transfer of small plasmids achieves higher efficiencies and the three streamlined PGC-containing ECRs still appear to confer the ability to perform photosynthesis (Kreiss *et al.*, 1999; Chan *et al.*, 2002). The acquisition of streamlined PGC-containing ECRs might enable strains to obtain new lifestyles at low costs, thereby providing advantages under certain environmental conditions (Petersen *et al.*, 2013). The other large PGC-containing ECRs also encoded proteins with various non-photosynthetic functions. For example, the PGC-containing chromid-like elements of *N. ignava* DSM16309 comprises 11 co-localized genes that enable sulfur oxidation. Furthermore, most of these large ECRs were classified as chromid-like ECRs. Consequently, these PGC-containing ECRs could preferentially be maintained in bacterial hosts, rather than be transferred among hosts. Notably, PGCs carried by both plasmid-like and chromid-like ECRs have been suggested to be genomically stable because most exPGCs have been inserted by their corresponding ECR replication modules (Petersen *et al.*, 2012).

Comparison of the GC content of the bacterial genomes, exPGCs, and PGC-containing ECRs did not reveal significant differences for any of the thirteen *Roseobacter* clade strains (Fig. S4). Thus, the transfer of these PGC-containing ECRs into bacteria likely occurred during very distant evolutionary events, or otherwise only between closely related species (Lassalle *et al.*, 2015).

### A scenario to explain the evolution of AAPB exPGCs in the *Roseobacter* clade

A previous scenario was suggested to explain the the evolution of exPGCs in *Roseobacter* clade organisms, wherein a chromosomal PGC superoperon was transferred into an ECR, followed by integration of replication origin genes into the ECR(Petersen *et al.*, 2012; Petersen *et al.*, 2013). Our analyses validate this explanation, and further present a more detailed scenario (Fig. 5) to explain the transfer of PGCs within the *Roseobacter* clade after analyzing genomic and evolutionary characteristics of thirteen exPGCs in this group (Petersen *et al.*, 2012). In this revised scenario, PGCs were first initially translocated from chromosomes to ECRs, as represented by the superoperon structure of exPGCs from *J. faecimaris* DSM1004020, *J. pohangensis* DSM19073, and *J.donghaensis* CECT 7802. Three subsequent transfer possibilities are present for exPGCs: (i) exPGCs reintegrated into chromosomes and became cPGCs as observed for most phototrophic *Roseobacter* clade strains; (ii) PGC-containing ECRs were lost from strains, such that bacteria became heterotrophs; or (iii) exPGCs carried by ECRs were subject to further recombination and became stable within ECRs. A remarkable characteristic of the majority of exPGCs is the insertion of an ECR replication module within the PGC as a result of a series of recombination events. Such an event could have helped ensure the stability of exPGCs (Petersen *et al.*, 2012).

**Fig. 5.**
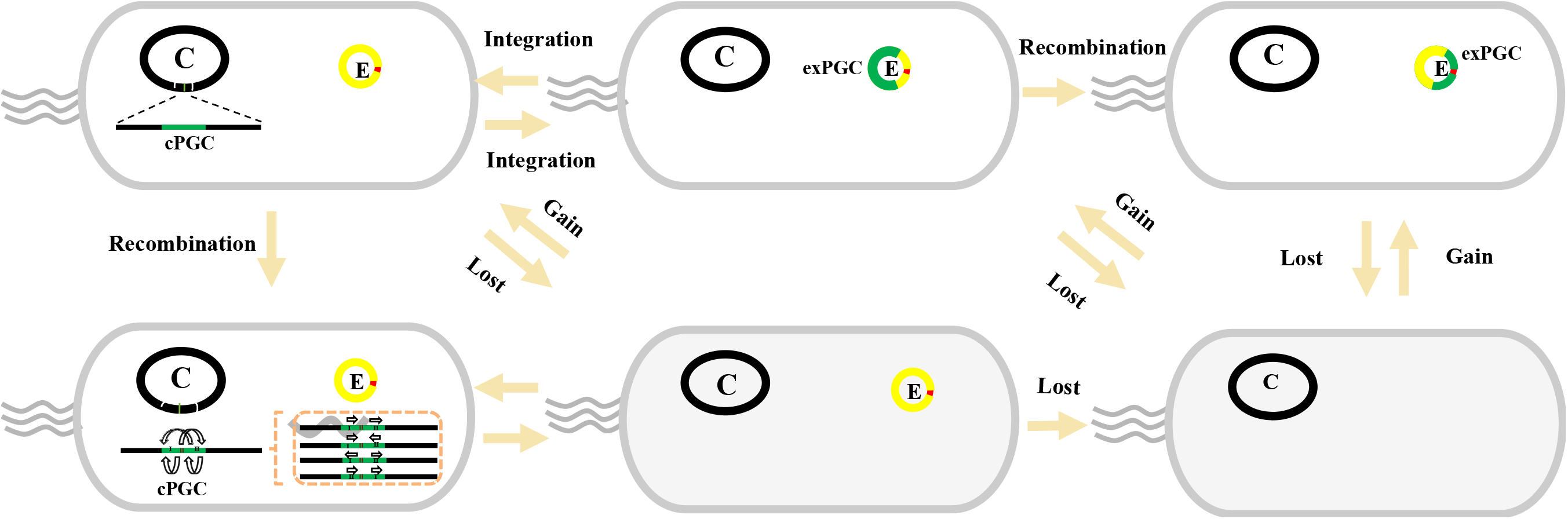
Scenarios to explain the transfer and evolution of exPGCs within *Roseobacter* clade species. PGCs are indicated in green and PGC-containing ECR origins of replication are indicated in red. Photosynthetic bacteria are indicated in light red, while non-photosynthetic bacteria are shown in light grey. C: chromosome, E: PGC-containing ECR.

The patchy distribution of AAPB within the *Roseobacter* clade has been explained by two evolutionary models that either invoke the loss or gain of PGCs (Yurkov and Csotonyi, 2009a; Yurkov and Hughes, 2017; Beatty, 2002). Given that ECRs play a critical role in the loss or gain of PGCs during the evolutionary history of photosynthesis, the patchy distribution of AAPB within the *Roseobacter* clade can be plausibly explained by ECR-mediated mechanisms. Until now, exPGCs account for ~20% of all PGCs in the currently available genomes of *Roseobacter* clade strains (Table S2), highlighting their prevalence in these organisms. It is likely that additional exPGCs carried by other strains will be identified with further generation of new bacterial genome sequences. Moreover, we suggest that the gain and loss of PGCs, as mediated by chromosomes and especially ECRs, result in the patchy distribution of AAPB within the *Roseobacter* clade.

In the present study, the genomic characteristics and evolution of thirteen PGCs carried by ECRs were analyzed. The coexistence of two different photosynthetic reaction complex types within strains of the same genera provided clear evidence of the horizontal transfer of PGCs mediated by ECR. Analysis of PGC-containing plasmid-like and chromids-like ECRs indicated that exPGCs could stably exist in bacteria after transfer, highlighting the importance of photosynthetic metabolisms carried by ECRs for some bacteria. Furthermore, these analyses indicated that the gain or loss of PGCs, as mediated by ECRs, contributes to the patchy distribution of photosynthetic capacity within the *Roseobacter* clade.

## Materials and methods

### Strain isolation

*Tateymaria* sp. syn59 and *Shimia* sp. wx04 were isolated from the South China Sea in April 2016 using oligotrophic medium F/2 plates (Guillard and Ryther, 2011), followed by transfer onto rich organic liquid medium (Difco Marine Broth 2216, American) for further isolation and cultivation. All cultures were incubated at 28°C with shaking at 200 rpm in the dark. Genomic DNA from the two strains was extracted using a Takara MiniBEST Universal Genomic DNA Extraction Kit (Japan).

### Genome sequencing, assembly, and annotation

The genomes of *Tateymaria* sp. syn59 and *Shimia* sp. wx04 were sequenced on the Illumina MiSeq platform (Illumina, USA). Specifically, 2 x 250 bp paired-end read sequencing was conducted, followed by read assembly using the velvet program (version 2.8) (Zerbino and Birney, 2008). Prediction and annotation of open reading frames (ORFs) was conducted using the Rapid Annotation using Subsystems Technology (RAST) platform. Further, the annotation of plasmids and exPGCs were validated by BLASTP searches against the National Center for Biotechnology Information (NCBI, https://blast.ncbi.nlm.nih.gov/Blast.cgi) non-redundant (nr) protein database.

### Retrieval of AAPB genomes from GenBank

Genome sequence data for the other eleven *Roseobacter* clade strains was obtained from NCBI including for *Tateyamaria omphalii* DOK1-4 (CP019312); *Tateyamaria* sp. ANG-S1 (JWLL00000000); *Roseobacter litoralis* Och 149 (CP002623); *Oceanicola* sp. HL-35 (JAFT00000000.1); *Nereida ignava* DSM 16309 (CVPC00000000); *Sulfitobacter noctilucicola* KCTC 32123 (JASD00000000.1); *Sulfitobacter guttiformis* KCTC 32187 (JASG00000000); *Sulfitobacter* sp. AM1-D1 (CP018076.1); *Jannaschia donghaensis* CECT 7802 (CXSU00000000); *Jannaschia faecimaris* DSM 10004020 (FNPX00000000.1); and *Jannaschia pohangensis* DSM 19073 (FORA00000000).

### Phylogenetic analysis

Complete 16S rRNA gene sequences were extracted from the whole genome assemblies (WGA) using the Cluster program (series1999). To construct a phylogeny for the PGCs, the amino acid sequences of 29 conserved photosynthetic genes within PGCs *(bchI, bchD, bchO, tspO, crtC, crtD, crtF, bchC, bchX, bchY, bchZ, pufL, pufM, bchP, puCC, bchG, ppsR, bchF, bchN, bchB, bchH, bchL, bchM, IhaA, puhA, puhB, puhC, ascF,* and *puhE)* (Table S3) were retrieved from the query genomes and then individually aligned using ClustalW, as implemented in the BioEdit program (Hall *et al.).* Phylogenetic analyses were conducted using RAxML 8.02 (Stamatakis, 2014) and maximum likelihood (ML) methods. The robustness of tree topologies were evaluated using bootstrap analysis with 100 replicates. Final trees were then visualized using the Interactive Tree of Life viewer and MEGA version 7.0 (Kumar *et al.*, 2016; Letunic and Bork, 2016).

## Acknowledgements

We are grateful to J. Thomas Beatty for helpful discussion and constructive suggestions. This work was supported by the National Key Research Programs (2018YFA0605800), the National Natural Science Foundation of China (NSFC) project (41776145, 41876150 and 91751207), and Natural Science Foundation of Fujian Province of China (2018J05072). All authors have no conflict of interest.

